# Live dynamics of induced cell-cell fusion between mitotic and interphasic cells

**DOI:** 10.64898/2026.01.27.700572

**Authors:** Olga Afonso, Daniel Feliciano, Jennifer Lippincott-Schwartz

## Abstract

The cell cycle is tightly regulated by checkpoint mechanisms that ensure faithful duplication and segregation of the genome. Here, we induced cell-cell fusion between mitotic and interphase cells to study how nuclei from different cell cycle stages behave in a shared cytoplasm. We found that mitosis is a dominant cell cycle state: the mitotic cytoplasm can drive interphase nuclei into mitosis, whereas, in high ratios of interphase versus mitotic nuclei, fusion forced mitotic nuclei to exit mitosis. Both outcomes represent checkpoint override events with impactful consequences. Interphase nuclei forced into mitosis form aberrant mitotic spindles, show partially condensed DNA and ultimately undergo mitotic catastrophe. Conversely, forced mitotic exit resulted in reformation of nuclear envelope membranes around condensed chromosomes, forming nuclei with a defective nuclear import machinery. Altogether, cell-cell fusion revealed an unexpected plasticity in cell cycle control and highlight cell-cell fusion experiments as a powerful experimental system to study how competing cytoplasmic states are integrated in a shared cytoplasm.

## Introduction

Cell division allows the propagation of unicellular species and the generation of tissues that form a multicellular organism. The cell cycle machinery, driven by the Cyclin/Cdk complexes, ensures a specific order of events essential for faithful transmission of the genetic material. Cyclins are expressed in an oscillatory pattern and produced at specific time points throughout the cell cycle (Evans *et al*, 1983). Regulation of cyclin expression, together with its Cdk partner, promotes the transition from one cell cycle stage to another through a series of signaling cascades known as checkpoints (Hartwell & Weinert, 1989). These not only define the order of cell cycle events but also “check” if a process is completed correctly, and if so, allow progression to the next stage. Different Cyclin/Cdk complexes drive different cell cycle processes. For example, Cyclin A and E associated with Cdk2 promote entry into S phase and DNA replication (Malumbres, 2014); progression to mitosis is triggered by increasing levels of Cyclin A and B bound to Cdk1 (Gavet & Pines, 2010b), and finally degradation of mitotic cyclins by the anaphase-promoting complex (APC/C) bound to Cdc20 or Cdh1 promotes nuclear envelope reformation around decondensing chromosomes and return to the interphase state (Peters, 2006), completing one cell cycle. Currently, nearly 20 Cdks and 30 Cyclins have been identified in mammals with different functions in cell cycle control (Malumbres, 2014).

Cell-cell fusion experiments with cells in different cell cycle stages were pioneered in the 70’s and 80’s (Ghosh *et al*, 1978; Ikeuchi & Sandberg, 1970; Johnson & Rao, 1970; Kato & Sandberg, 1968; Matsui *et al*, 1972) before Cdks and cyclins were identified. The majority of these experiments showed that interphase cells were efficiently induced to enter mitosis after fusion with one or more mitotic cells (Hanks & Rao, 1980; Johnson & Rao, 1970, 1971; Johnson *et al*, 1970), but also, mitotic and interphase cell-cell fusion could lead to premature mitotic exit, with metaphase-like chromosomes surrounded by nuclear envelope membranes (Ghosh & Paweletz, 1987a, b; Ikeuchi *et al*, 1971). These experiments provided important insights into cell cycle control and demonstrated the suitability of cell-cell fusion experiments in studying cell cycle transitions.

Under physiological conditions cell-cell fusion occurs spontaneously and is essential for the development of different tissues such as muscle, bone and placenta (Ogle *et al*, 2005). It has also been implicated in cancer expansion through cell-cell fusion events between cancer cells and stem cells (Bjerkvig *et al*, 2005). However, it is not known whether cell-cell fusion events occur only between cells at the same cell cycle stage or if it can occur between cells at different stages.

Here we revisited these classic experiments by combining the viral VSV-G protein to induce fast cell-cell fusion between mitotic and interphase cells with state-of-the-art live-cell microscopy technology to study the outcomes of the formed syncytium. We found that, in the syncytium, the mitotic cytoplasm is dominant and capable of inducing mitotic entry in up to six neighboring nuclei, forming mitotic syncytia that are not viable and frequently undergo mitotic catastrophe. In the presence of seven or more interphase nuclei, the most frequent outcome was forced mitotic exit of mitotic nuclei, where nuclear envelope membranes reformed around condensed chromosomes. Our data raises the question of how cell-cell fusion is regulated under physiological conditions to avoid the generation of failed syncytia that can be deleterious to the tissue.

## Results and Discussion

### MITOTIC entry in the syncytium depends on proximity to mitotic chromosomes

Cell-cell fusion assays performed in the 70’s and 80’s relied primally on electron microcopy analysis of fixed samples and therefore captured the final outcome of fusion events. To directly examine how the tug-of-war between interphase and mitotic cells is resolved over time, we used high-resolution live-cell imaging microscopy of fusion events in asynchronously growing cell cultures. Briefly, cell-cell fusion was induced by transient expression of the vesicular stomatitis virus G-protein (VSV-G), which induces nearly immediate cell-cell fusion after an acute low pH pulse (Feliciano *et al*, 2018). To identify fused cells, we transiently co-transfected a blue fluorescent protein (BFP): when cells fuse and the cytoplasm mixes, the cytoplasmatic BFP signal is redistributed and becomes dispersed into all cells that form the syncytium (Figure 1A). In this study, we used LLC-PK1 cells (pig kidney cells) which are flat cells, ideal for high resolution live imaging, and tolerate formation of large syncytia without detaching. To identify nuclei and cell boundaries, we used a cell line expressing a nuclear marker (H2B-mCherry) and a reticulum endoplasmic marker (ER3-emerald). Note that, despite being a stable cell line, not all cells showed the expression of the nuclear marker (example in Figure 1A).

**Figure 1.**
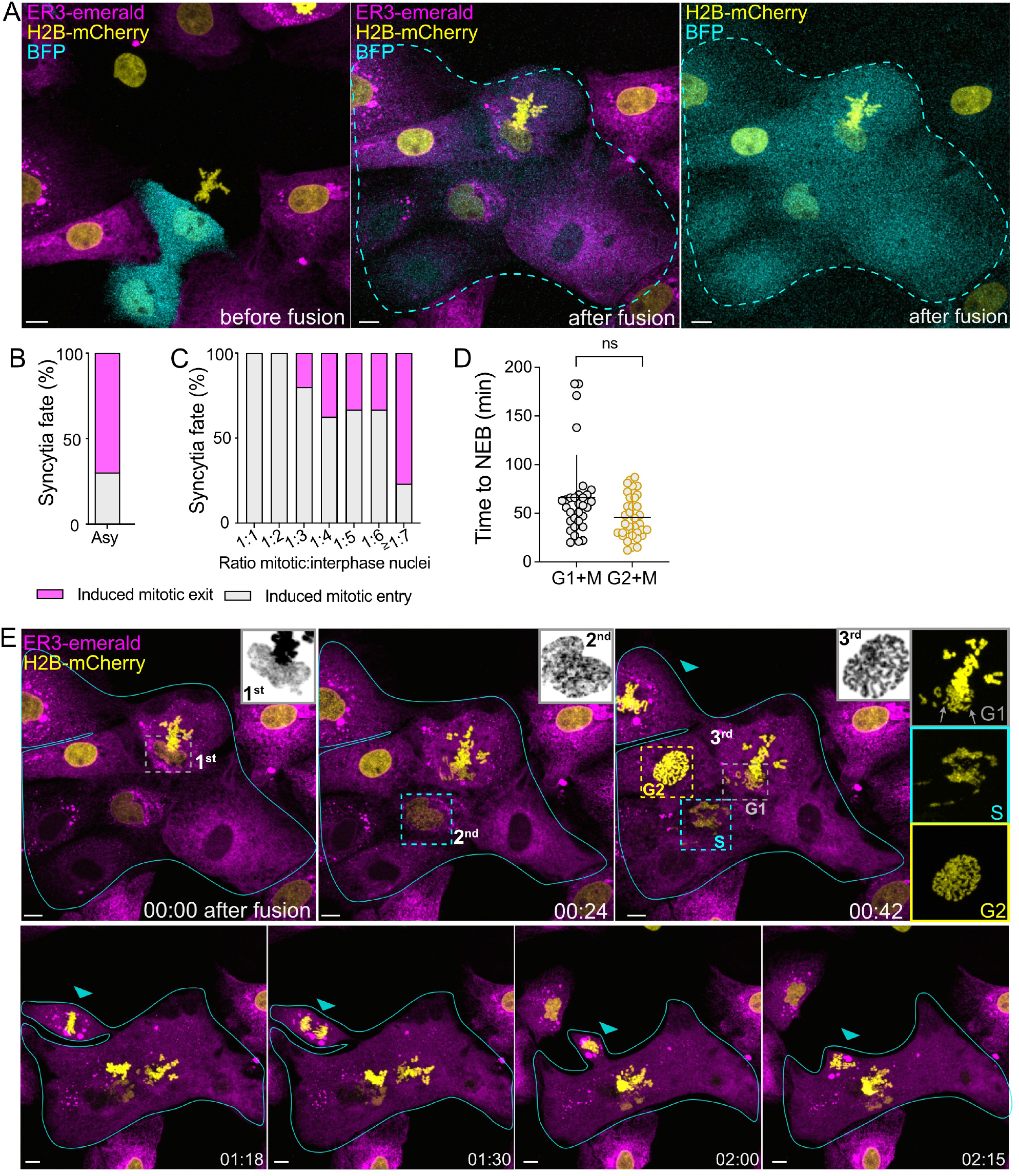
Ratio of fused mitotic and interphase nuclei determines syncytium fate. (A) Example of induced mitotic entry after cell-cell fusion in an asynchronous culture of LLC-PK1 cells expressing H2B-mCherry/ER3-emerald and transiently transfected with Blue Fluorescent Protein (BFP), to detect fusion. Note that the acquisition window was re-adjusted from the “before fusion” to “after fusion” images to capture the entire syncytium. (B) Percentage of syncytia where mitotic entry or mitotic exit was observed after induced cell-cell fusion in asynchronous cell cultures (n=10 syncytia). (C) Mitotic:interphase nuclei ratio analysis and its impact on the fate of the syncytium from a pool of asynchronous (n=10), G1+M (n=23) and G2+M (n=19) syncytia. (D) Time from induced fusion until nuclear envelope breakdown (NEB) in G1+M (n=29 nuclei) and G2+M (n=37 nuclei) enriched cultures. Statistics, non-parametric Mann-Whitney test. (E) Time-lapse of the same syncytium as in (A) showing asynchronous entry into mitosis (1^st^, 2^nd^ and 3^rd^ cells) and the different cell cycle stages based on the condensed chromatin structure: single chromatids in G1 nuclei (grey squares and arrows), unstructured chromatin in S-phase nuclei (cyan squares) and condensed chromosomes with two chromatids in G2 nuclei (yellow squares). Note that the top left corner cell is fused by a narrow bridge, enters mitosis independently (blue arrowhead) and joins the syncytium after anaphase (“02:00” time point). Time is in h:min. Scale bar is 10μm.

Live imaging of fusion events between mitotic and interphase cells showed only two possible outcomes: either all nuclei entered or exited mitosis (Figure 1B). Interestingly, we found that a single mitotic nucleus was sufficient to drive mitotic entry in up to six interphase nuclei within the same syncytium, suggesting that the mitotic cytoplasm is dominant over the interphase cytoplasm. Only at a ratio of seven or more interphase cells per mitotic cell, forced mitotic exit of mitotic cells became the most frequent outcome (Figure 1C).

We first focused on syncytia in which all nuclei entered mitosis (mitotic syncytia). Although cytoplasm mixing after fusion can be achieved rapidly within two minutes (Feliciano *et al*., 2018), nuclei within the same syncytium entered mitosis at different timings, as determined by chromatin condensation (Figure 1E and Figure 2A). Also, using the morphology of condensed chromatin within a syncytium, we could distinguish nuclei from different cell cycle stages: G1 nuclei with condensed single chromatids, S-phase nuclei, with unstructured condensed DNA and G2 nuclei, with condensed chromosomes with two chromatids (Figure 1E, grey, cyan and yellow squares, respectively, and Figure S1).

**Figure 2-.**
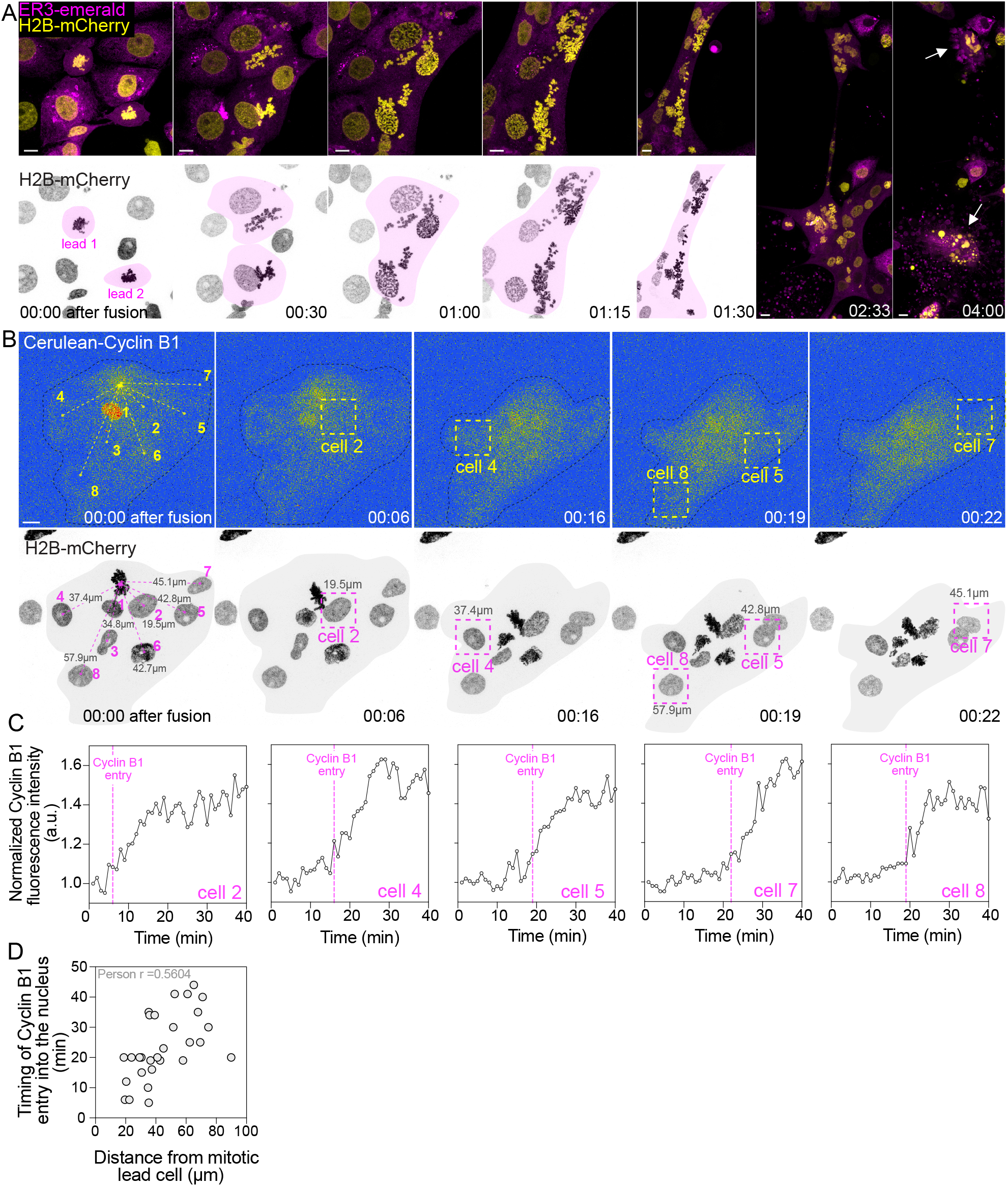
Mitotic entry in the syncytium is dependent on the close proximity to mitotic chromosomes. (A) Time-lapse of a mitotic syncytium in LLC-PK1 cells expressing H2B-mCherry/ER3-emerald. Magenta shadowed areas highlight the progression of induced mitotic entry over time. Not all cells enter mitosis at the same time. (B) Time-lapse of a mitotic syncytium expressing Cerulean-Cyclin B1 and H2B-mCherry. Cells are numbered by increasing distance from the lead mitotic cell and their distances are shown. Dashed black line and grey shadow show syncytium area and magenta squares highlight progressive mitotic entry of cells in the syncytium. Time is in h:min. Scale bar is 10μm. (C) Normalized Cyclin B1 fluorescence intensity in several nuclei of the mitotic syncytium in (B). Cyclin B1 accumulates first on nuclei close to the mitotic lead cell. Cyclin B1 intensity values are normalized to the first frame after fusion. (D) Positive correlation between time of Cyclin B1 entry in the nucleus and their distance from the lead mitotic cell.

We therefore hypothesized that the out of phase mitotic entry reflected the initial cell cycle stage of each nucleus. To test this, we induced cell-cell fusion between mitotic cells (M) and cells arrested at either G1/S (G1/S+M fusion) or at G2 (G2+M fusion). In both conditions we observed the asynchronous pattern of mitotic entry but no statistical differences in the timing of mitotic entry between the G1/S+M and G2+M fusions (Figure 1D). These results suggest that asynchronous mitotic entry within a syncytium is not determined by the initial cell cycle state of the nucleus.

Instead, we observed that the interphase nuclei closest to mitotic cells present at the moment of fusion entered mitosis first (Figure 2A). In most mitotic syncytia, one or two initial mitotic cells (lead cells) triggered mitotic entry in the neighboring interphase nuclei. These in turn, induced mitotic entry in additional adjacent nuclei, resulting in a wave-like propagation of induced mitotic entry across the syncytium (Figure 2A, magenta shadowed area). Importantly, within 30-60min after fusion, nuclei have moved in the syncytium (Figure 2A, compare time “00:00” *vs* “01:30”) and their positions relative to the initial mitotic lead cells can no longer be tracked. Therefore, the wave-like pattern of mitotic entry is better detected within the first 30-60min after fusion. Out of ten mitotic syncytia analyzed, four underwent mitotic catastrophe (Figure 2A, last time point, white arrows) while the others remained in mitosis until the end of acquisition (5 hours).

It is well established that mitotic entry is triggered by the presence of active Cdk1, through binding to Cyclin B1 (Nurse, 1990). CyclinB1/Cdk1 is distributed in the cytoplasm and enters the nucleus ∼30 minutes before nuclear envelope breakdown (NEB) (Gavet & Pines, 2010a). To verify the distribution of CyclinB1/Cdk1 after cell-cell fusion, LLC-PK1 cells stably expressing H2B-mCherry were transiently transfected with cerulean-Cyclin B1. Interestingly, while Cyclin B1 was homogenously distributed in the cytoplasm quickly after fusion, Cyclin B1 entry into the nucleus was asynchronous (Figure 2B and C), happening on average 10 minutes before DNA condensation. Importantly, timing of Cyclin B1 entry into the nucleus correlated with their distance to the lead mitotic cell, at the moment of fusion (Figure 2D), suggesting that mitotic entry within a shared cytoplasm may not be only driven by the homogeneous cytoplasmic pool of Cyclin B1/Cdk1. Instead, mitotic chromosomes seem to act as local signaling hubs that accelerate mitotic entry in neighboring interphase nuclei, resulting in a distance-dependent, wave-like propagation of mitosis across the syncytium. Indeed, Cyclin B1/Cdk1 are enriched on mitotic chromosomes (Clute & Pines, 1999), suggesting the presence of a gradient of activity away from the chromosomes. Using the Cdk1 FRET sensor (Gavet & Pines, 2010b) with high resolution live imaging would clarify this hypothesis.

### Mitotic structures assembled on condensed G1/S phase DNA are defective

During embryonic development, the one-cell stage embryo is a very large cell (up to 1mm in length in *Xenopus laevis*) that goes through successive cell division cycles generating smaller and smaller cells. Under these conditions, the size of the mitotic spindle has two properties: it has a maximum length in the largest cells (∼60μm in *Xenopus*) and, as cells become smaller, spindle length is maintained proportional to cell size, i.e., the spindle length scales with cell size (Rieckhoff *et al*, 2020; Wuhr *et al*, 2008). Interestingly, mitotic syncytia ranged several orders of magnitude in size, with the largest syncytia being within the same size range as a zebrafish embryo (∼0.5mm in length and can be detected by naked eye). Thus, we tested if in the mitotic syncytia, spindle size would also scale with syncytia size.

We found that the average spindle length per syncytium did not correlate with syncytium area, but the total number of spindles did (Figure 3B and C). Indeed, within a syncytium, the DNA was often organized around multiple bipolar spindles, multipolar spindles and/or arrays of connected bipolar spindles, but never a single large spindle (Figure 3A and F). This agrees with previous *in vitro* studies where spindles assembled on DNA coated bead strings showed a maximum spindle size regardless of string length (Gaetz *et al*, 2006). Thus, in contrast to embryonic development, after cell-cell fusion, in somatic cells each spindle behaved as an independent identity regardless of syncytium size or DNA amount. In order to scale, there must be a mechanism to sense cell size and, potentially, the size of mitotic syncytia was too large for the spindle to reach the boundaries and sense the new syncytium size.

**Figure 3-.**
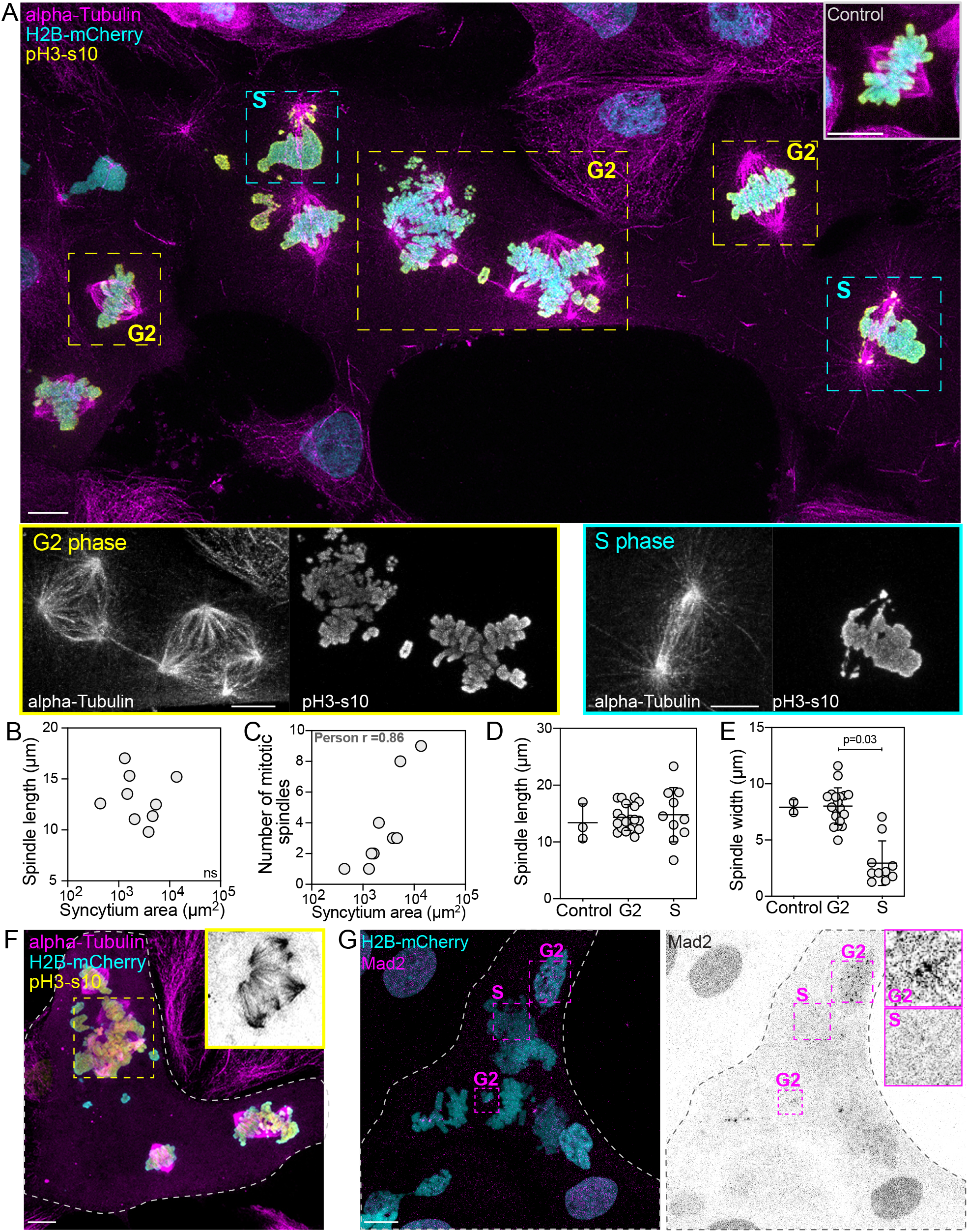
Mitotic spindles assembled on condensed G1/S DNA are defective. (A) Example of a mitotic syncytium stained for microtubules (magenta), chromosomes (cyan) and phosphorylated pH3s10 (yellow). G2 chromosomes interact with microtubules and form bipolar and multipolar spindles (yellow dashed squares). S-phase chromosomes assemble defective spindles (cyan dashed squares). Up right corner, control mitotic cell for comparison. (B) Correlation between average spindle length in a syncytium and respective syncytium area (n=9 syncytia). No significant correlation was observed. (C) Positive correlation between the total number of mitotic spindles in a syncytium and respective syncytium area (n=9 syncytia). (D and E) Quantification of spindle length and spindle width, respectively, in control (n=3), G2 (n=19) and S-phase (n=10) bipolar spindles in syncytia. (F) Example of a mitotic syncytium where an array of mitotic spindles was formed around different types of condensed DNA. (G) Example of a mitotic syncytium stained for Mad2 (magenta) and chromosomes (cyan). Mitotic chromosomes and early prometaphase nuclei (“G2”) show positive Mad2 at kinetochores, however the signal is not detected in S-phase DNA (“S”). Scale bar is 10μm.

Mitotic syncytia are formed by nuclei from different cell cycle stages. Thus, we were interested in understanding how the mitotic spindle is organized on different forms of condensed DNA. G2-phase chromosomes showed stable attachments to microtubules, forming bipolar spindles like control cells, but also multipolar spindles (Figure 3A, yellow dashed squares and control cell). Spindle length and width of “G2 spindles” was comparable to that of control cells (Figure 3D and E). In contrast, S-phase DNA showed spindles with few microtubules interacting with the DNA (Figure 3A, cyan dashed squares). While spindle length was not significantly different from control cells, spindle width was (Figure 3D and E), reflecting the low microtubule density of S-phase spindles.

At the basis of mitotic spindle assembly is the interaction between microtubules and kinetochores, a multi-subunit protein complex that is established at the centromeric region of each sister chromatid (Musacchio & Desai, 2017). The absence of stable kinetochore-microtubule interactions leads to less robust spindles without visible microtubule bundles (DeLuca *et al*, 2005; DeLuca *et al*, 2002), similar to the spindles observed on S-phase chromosomes. To test if S-phase DNA assembles functional kinetochores, we tested the localization of the Mad2 protein in both S and G2 DNA. Mad2 belongs to the Spindle Assembly Checkpoint machinery (SAC) and signals for the presence of unattached kinetochores: if kinetochores are not bound to microtubules Mad2 is present at kinetochores and delays progression into anaphase (Musacchio, 2015). In the syncytium, duplicated unaligned mitotic chromosomes and early prometaphase nuclei show Mad2 recruitment (Figure 3G, “G2” dashed squares), while S-phase condensed DNA did not show Mad2 enrichment (Figure 3G, “S” dashed squares).

Our data suggests that S phase kinetochores are not properly assembled, impairing Mad2 loading. Interestingly, S-phase DNA in mitotic syncytia resembles the DNA organization described in cells that undergo mitosis with unreplicated genomes (MUGs) (O’Connell *et al*, 2009). However, in MUGs, kinetochores are functional as they recruit checkpoint proteins such as Mad2 (Johnson *et al*, 2008) and interact with microtubules to establish a bipolar spindle (O’Connell *et al*., 2009). These differences could be related with the fast induced mitotic entry of interphase nuclei that does not provide time for the assembly of a functional kinetochore.

All together, these results show that although cells in any stage of the cell cycle can rapidly switch into a mitotic state, non-replicated DNA cannot assemble proper kinetochores or bipolar spindles, resulting a syncytium that either becomes arrested in mitosis or undergoes mitotic catastrophe.

### Cell-cell fusion induced mitotic exit is independent of anaphase events

One of the outcomes of induced cell-cell fusion between mitotic and interphase cells is a switch from the mitotic to an interphase-like state. In these interphase syncytia, nuclear envelope membranes reformed around apparently condensed chromosomes from mitotic cells (Figure 4A), in agreement with previous reports of cell-cell fusion experiments (Ikeuchi *et al*., 1971). To test this, we quantified H2B-mCherry intensity over time in anaphase control cells and in mitotic fused cells undergoing mitotic exit. Chromosomes reach their maximal compaction during anaphase, a process dependent on Aurora B kinase (Mora-Bermudez *et al*, 2007). Accordingly, control cells showed a maximum of H2B intensity approximately 14 minutes after anaphase onset and the intensity reduced from that moment onwards (Figure 4B). In mitotic cells forced to exit mitosis, H2B intensity was largely constant, from the moment of fusion until up to 60 min after fusion (Figure 4B). Additionally, the nucleus of control cells achieved a sphere-like shape 12 min after anaphase onset, but mitotic fused cells it did not change their shape significantly over the time window of image acquisition (Figure 4A and C). These results suggest that DNA decondensation does not occur during exit of mitotic fused cells.

**Figure 4.**
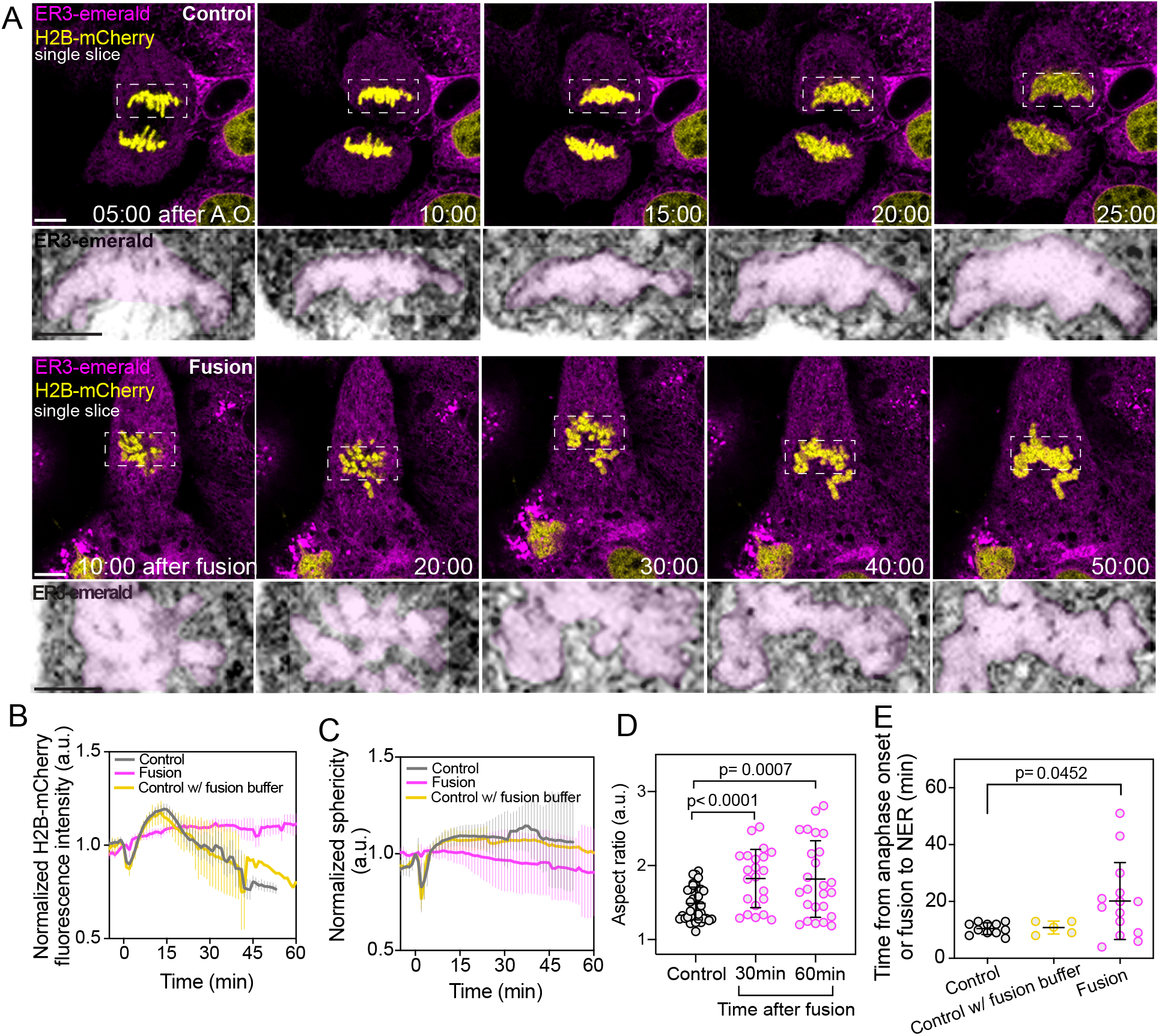
Induced mitotic exit after cell-cell fusion leads to nuclear envelope reformation without anaphase events. (A) Time lapse of a control cell progressing through anaphase and mitotic fused cells undergoing premature mitotic exit. Chromosomes are in yellow and nuclear envelope in magenta. In premature mitotic exit, the nuclear envelope reforms without chromosome separation. Time is in min:sec. Scale bar is 10μm. (B and C) Quantification of H2B-mCherry fluorescence intensity and sphericity, respectively, in control cells without (n=10 cells) and with (n=4 cells) fusion buffer and in mitotic fused cells (n=8 cells). (D) Quantification of nucleus aspect ratio, showing that nucleus shape after induced mitotic exit does not achieve the round shape characteristic of interphase control nuclei. Control, n=20 cells; 30 min, n=22 cells; 60 min, n=25 cells. Statistics, parametric t-test. (E) Quantification of anaphase duration in control cells without (n=11 cells) and with (n=5 cells) fusion buffer, or time from fusion to nuclear envelope reformation in mitotic fused cells (n=14 cells). Statistics, non-parametric Mann-Whitney test.

The abnormal nucleus shape after induced mitotic exit was clear in fixed cell analysis, with a significantly higher aspect ratio of nuclei induced to exit mitosis (Figure 4D). As mitotic exit in mitotic fused cells showed dissimilarities from mitotic exit in control cells, we determined the timing of NER by analyzing the accumulation of the reticulum endoplasmic marker ER3-emerald on mitotic chromosomes. Using as reference the moment of fusion, membranes were identified around chromosomes of mitotic fused cells on average 20 min after fusion, showing a slower kinetics of mitotic exit than control cells (Figure 4E).

Our data suggests that, during premature mitotic exit of mitotic cells, chromosomes remain condensed however nuclear envelope membranes reform around them. This condition is unique as in unperturbed anaphase cells, DNA decondensation and nuclear envelope occur at the same time and are regulated by the balanced activity of Aurora B kinase and counteracting phosphatases PP1 and PP2A (Afonso *et al*, 2014). Since these two events seem indissociable, the general assumption in the field is that DNA decondensation is required for NER. Thus, we used the cell-cell fusion assay to further investigate if the condensed chromosomes were still in a “mitotic state”.

We observed that during mitotic exit after cell-cell fusion, sister chromatids do not separate neither move to opposite poles, and fused cells do not undergo cytokinesis (Figure 4A and 5C). These events occur in control anaphase cells due to activation of the E3 ubiquitin ligase APC/C by the co-activator Cdc20. Active APC/C-Cdc20 triggers Cyclin B1 and Securin degradation, subsequent Cdk1 down regulation and Aurora B re-localization from the centromeres to the spindle midzone (Pines, 2006). Thus, fused mitotic cells seem to have skipped anaphase and proceeded immediately to telophase, without APC/C-Cdc20 activation.

As APC/C-Cdc20 activation is dependent on spindle assembly checkpoint silencing, we tested the localization of checkpoint proteins at kinetochores after nuclear envelope reassembly on mitotic fused cells. In control cells, Mad2 localized to unattached kinetochores during prometaphase and was absent in anaphase and telophase (Figure 5A). In mitotic cells where cell-cell fusion induced mitotic exit, Mad2 was clearly visible at kinetochores of mitotic chromosomes even after nuclear envelope reassembly (Figure 5A). If APC/C-Cdc20 is not activated during induced mitotic exit, we would expect high levels of Cyclin B1 in those cells. In agreement, in the rare event of mitotic exit after expression of exogenous Cyclin B1, since the Cyclin B1 overexpression further potentiates mitotic syncytia, we observed that mitotic cells exited mitosis in the presence of high levels of Cyclin B1 on the chromosomes (Figure 5B).

**Figure 5-.**
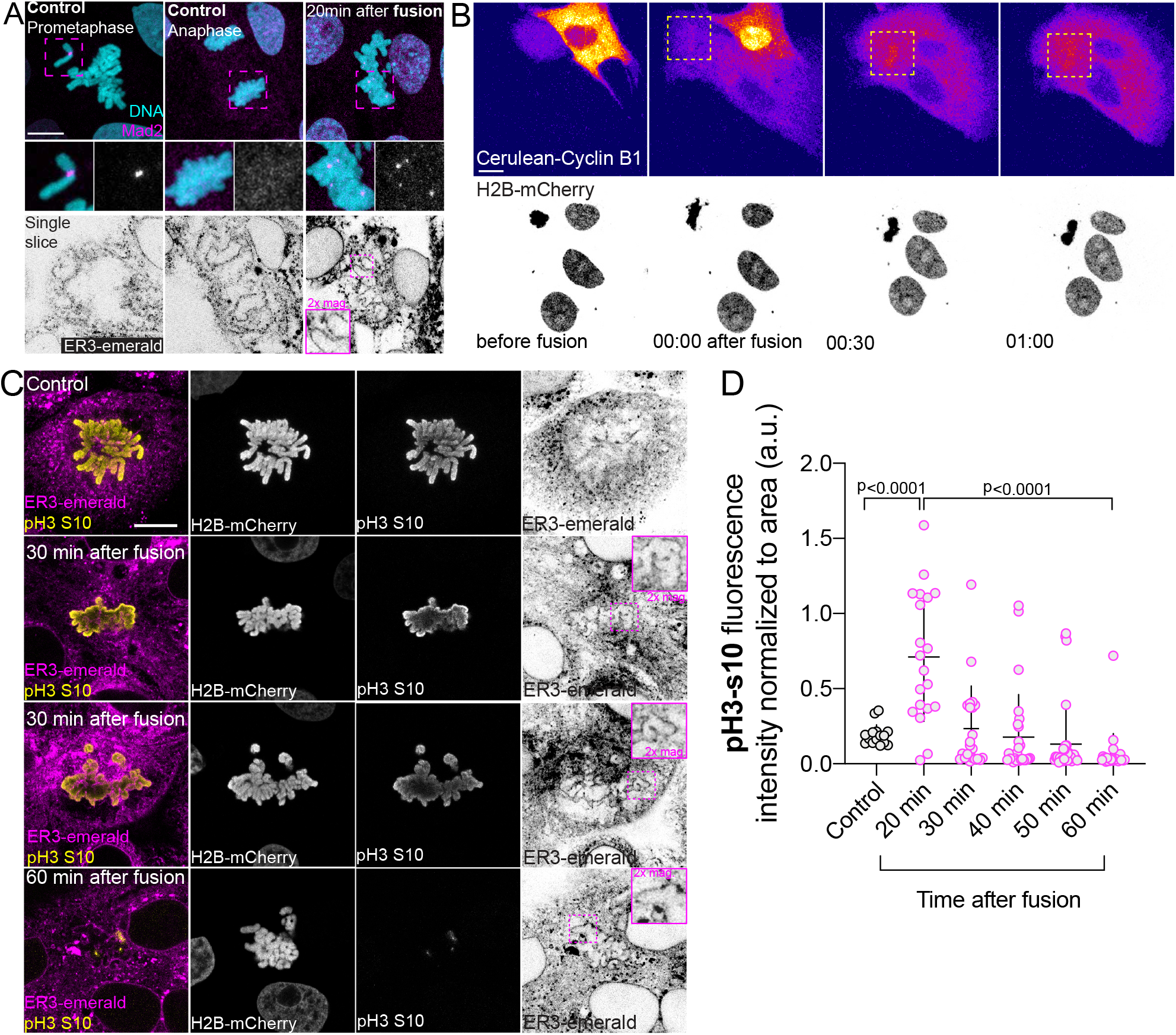
Nuclear envelope membranes are prematurely reformed around mitotic chromosomes. (A) Comparison of Mad2 localization in control prometaphase and anaphase cells and mitotic fused cells undergoing premature mitotic exit. Mad2 can be detected at some kinetochores despite the presence of nuclear envelope membranes. (B) Time lapse of cerulean-Cyclin B1 transiently transfected in LLC-PK1 cells expressing H2B-mCherry after cell-cell fusion. Cyclin B1 was not degraded during induced mitotic exit of the mitotic cell. (C) Analysis of pH3-s10 in fixed prometaphase control cells and at different time points after cell-cell fusion. At 30 minutes after cell-cell fusion there is a mixed population of cells with high (second panel) and low (third panel) pH3-s10 levels, despite the presence of nuclear envelope membranes. At 60 minutes after cell-cell fusion, most cells show very low levels of pH3-s10. Magenta squares show magnification of nuclear envelope membranes around chromosomes. (D) Quantification of pH3-s10 in telophase control cells and at several time points after cell-cell fusion. Control, n=19 cells; 20 min, n=20 cells; 30 min, n=22 cells; 40 min, n=27 cells; 50 min, n=32 cells and 60 min, n=26 cells. Note that only 50 and 60 minutes after cell-cell fusion the levels of pH3-s10 become similar to control levels. Statistics, non-parametric Mann-Whitney test. Time is h:min. Scale bar is 10μm.

Finally, a well-established marker of mitotic exit is dephosphorylation of Histone H3 at Ser10. Histone H3 phosphorylation at Ser10 (pH3s10) is a result of the targeted action of Aurora B kinase which is counteracted by phosphatases during mitotic exit (Afonso *et al*., 2014; Fuller *et al*, 2008). Importantly, pH3s10 is associated with DNA condensation status (Giet & Glover, 2001; Sauve *et al*, 1999; Wei *et al*, 1998).

In control cells, pH3s10 showed the expected pattern: high levels in prometaphase and metaphase cells, telomere localization during anaphase and nearly absent in telophase, when the nuclear envelope starts to reform (Figure S2) (Afonso *et al*., 2014; Fuller *et al*., 2008; Hayashi-Takanaka *et al*, 2009; Van Hooser *et al*, 1998). Surprisingly, mitotic fused cells wrapped in nuclear envelope membranes showed high levels of pH3s10 that only decreased to control levels after 50 min after fusion (Figure 5C and D). This suggests that when nuclear envelope membranes reform, chromosomes are still highly phosphorylated and “in mitosis”. Using the Aurora B phosphorylation sensor (Afonso *et al*., 2014; Fuller *et al*., 2008) would allow us to understand if the phosphorylation levels remain high due to sustained kinase activity or inability of phosphatases to dephosphorylate substrates.

Altogether, our data shows that nuclear envelope membranes can reform around condensed and phosphorylated mitotic chromosomes uncoupling anaphase events, such as chromosome movement and DNA decondensation, from nuclear envelope reformation.

### Nuclear envelopes of mitotic fused cells are non-functional

A major question is whether the nuclear envelope that reforms on mitotic chromosomes is functional. Therefore, we tested the recruitment of other nuclear envelope components and functionality by looking at nuclear pore formation and nuclear import/export, respectively.

We used the monoclonal mab414 antibody that recognizes a pool of nucleoporins with phenylalanine-glycine (FG) repeat domains (Cronshaw *et al*, 2002; Davis & Blobel, 1986). While nuclear envelope membranes are detected around chromosomes at 20 minutes after fusion, these membranes lacked nuclear pore proteins and when present, nuclear pores were organized as large aggregates or patches instead of the continuous dot-like pattern observed in control telophase nuclei (Figure 6A and B). We also tested the recruitment of an inner nuclear membrane protein, lamin B receptor (LBR) in live cells. Live-imaging data of transiently expressed EGFP-LBR showed a slower recruitment dynamics in mitotic fused cells compared to control (Figure S3), similarly to what was observed for ER3 (compare Figure 4E and Figure S3C). Overall, our data shows that, while membranes can form around condensed chromosomes, they reform more slowly and fail to recruit nuclear pore proteins.

**Figure 6-.**
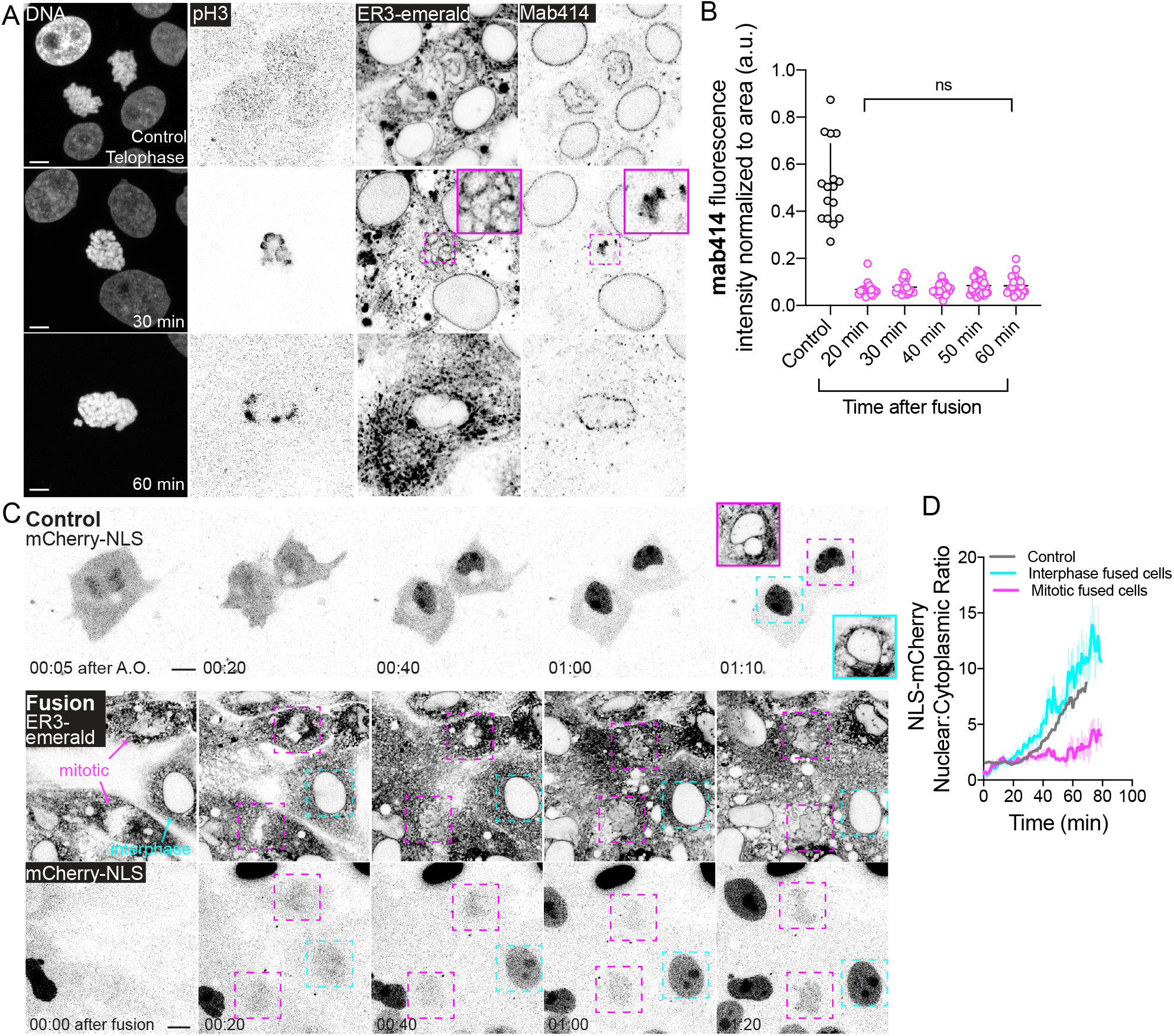
Nuclear envelopes of mitotic fused cells undergoing mitotic exit are non-functional. (A and B) Analysis of nuclear pore formation in control telophase cells and cells fixed at different time points after cell-cell fusion and respective fluorescence intensity quantification. Magenta square shows nuclear envelope membranes on condensed chromosomes but no nuclear pore protein accumulation. Control, n=19 cells; 20 min, n=20 cells; 30 min, n=22 cells; 40 min, n=27 cells; 50 min, n=32 cells and 60 min, n=26 cells. Statistics, non-parametric Mann-Whitney test. (C) Time-lapse of mCherry-NLS in control and mitotic fused cells shows efficient import of the NLS signal in control and fused interphase nucleus (cyan arrow), but not in mitotic fused nucleus (magenta arrows). Time is in h:min. Scale bar is 10μm. (D) Ratio of nuclear and cytoplasmic NLS signal of the examples shown in (C).

The lack of nuclear pore proteins suggests that nuclear import/export machinery does not occur. To test this, we performed a transient transfection of a nuclear localization signal (NLS) in cells stably expressing ER3-emerald. In control cells, NLS is imported into the nucleus during telophase (Figure 6C and D), showing that the nuclear pore import/export machinery is restored by the end of mitosis. In the syncytia, nuclei from cells that were in interphase before fusion showed an accumulation of NLS similar to what was observed for control cell exiting mitosis (Figure 6C and D). We reasoned that accumulation of NLS on interphase nuclei after fusion was a response to a new equilibrium of available NLS from the pool of fused cells that shared the same cytoplasm. In contrast to interphase nuclei, mitotic nuclei showed a slow accumulation of NLS and never reached the same levels as the interphasic neighbor nuclei (Figure 6C and D).

Altogether, the NLS and nuclear pore assembly data show that the nuclear envelope reformed on mitotic chromosomes is not functional. This can be either because the nuclear envelope has defects on the import machinery or membranes are not fully sealed and the NLS signal leaks to the cytoplasm. Interestingly, electron microscopy images of nuclei formed after fusion-induced mitotic exit show continuous nuclear envelope membranes at 40min after fusion (Ikeuchi *et al*., 1971). Therefore, nuclear envelope membranes can be recruited to mitotic chromosomes despite their condensation or phosphorylated state, however the nuclear pore machinery fails to assemble properly leading to a failure in nuclear import/export.

Interestingly, the results shown here on pH3s10 positive condensed chromosomes of mitotic fused cells are reminiscent of what has been reported on lagging chromosomes in human cells: lagging chromosomes remain pH3s10 positive for longer times than fully separated chromosomes (Afonso *et al*., 2014; Orr *et al*, 2021), they recruit nuclear envelope membranes (Liu *et al*, 2018) but fail to recruit nuclear pore proteins (de Castro *et al*, 2018; Liu *et al*., 2018; Orr *et al*., 2021). Lagging chromosomes have been shown to undergo chromothripsis and contribute to genomic instability. Therefore, it would be interesting to follow the fate of nuclei formed through forced mitotic exit - mitotic nuclei after cell-cell fusion are easily reproduced, can be obtained in large scale and are larger objects than lagging chromosomes, making this a useful system to study nuclear envelope reassembly on lagging DNA.

Taken together, our study reveals that cell-cell fusion between mitotic and interphase cells creates a dynamic tug-of-war between competing cell cycle states, with mitotic cytoplasm exerting dominance over interphase nuclei (Figure S4 and Figure 1B). We showed that in mitotic syncytia, interphase cells are induced to enter mitosis in a wave-like pattern that correlates with the physical distance to the nearest mitotic cell (Figure 2A) and nuclei with unreplicated DNA showed severe defects in spindle assembly and kinetochore function (Figure 3A). Conversely, premature mitotic exit resulted in the recruitment of nuclear envelope membranes to condensed phosphorylated (pH3s10 positive) chromosomes (Figure S4, Figure 4A and 5A), however, these nuclei lack nuclear pores and show impaired nuclear import function (Figure 6A and C). These results provide mechanistic insights into how cell cycle regulation is enforced or overridden in a shared cytoplasm and establish fused syncytia as a novel system to study the interplay between cytoplasmic cues, chromatin condensation state, and nuclear envelope reformation.

## Acknowledgments

This work initiated during the 2016 Physiology course edition under the supervision of J.L.S and D.F where O.A was a course participant. The project continued in J.L.S’s lab thanks to an MBL Post-Course research fellowship to O.A. We would like to thank the MBL for support during the course and post-course research. We also thank Cristopher Obara, Jonathan Nixon-Abell and members from J.L.S’s lab for support and discussions during project development. Finally, we thank Andrew S. Moore, Cristopher Obara, Marcos González-Gaitán and Ana Carvalho for critical reading of the manuscript.

## Data availability

Source data are provided with this paper. All other data supporting the findings of this study are available from the corresponding authors upon request.

## Competing interests

The authors declare no competing interests.

## Author contributions

O.A. performed experiments and analysis of experimental data with contributions from D. F. O.A., D.F. and J.L.S. conceived and designed experiments. O.A. prepared figures and manuscript with contributions from all authors.

## Methods

### Microscopy

Time-lapse microscopy and fixed cell analysis were performed in all experiments in a Zeiss LSM 880 confocal microscope, with acquisition every 3 or 1 minute. During acquisition cells were maintained in a temperature and CO_2_ incubated chamber.

### Cell culture

LLC-PK1 cells were cultured in DMEM supplemented with 10% FBS and grown in a 5% CO2 atmosphere at 37°C. For live-cell imaging experiments LLC-PK1 cells were plated at least 24h before the experiments in 2, 4 or 8-well Labteks at low concentration to avoid the formation of massive syncytia. Transfection with VSV-G and other plasmids was performed 10-12h before experiments.

### Immunofluorescence

Cells were fixed with 4% PFA, permeabilized with 0.3% Triton diluted in PBS and washed with PBS-Tween with 3 cycles of 5 min each. Primary mouse mab414 (1:1000; ab24609, Abcam), mouse anti-α-Tubulin (1:500, DM1A, sigma), mouse anti-Mad2 (1:100, sc-65492, Santa Cruz) and rabbit anti-pH3-s10 (1:200, 3377, Cell Signaling). The respective Alexa-Fluor secondary antibodies were used at 1:1000 dilution.

### Plasmids and transfections

All transient transfections were performed with Lipofectamine 3000 (Thermo Fisher Scientific) according to the manufacturer instructions. Cells were analyzed 10-12h after transfection with VSV-G. The EGFP-LBR plasmid has been previously described (Ellenberg *et al*, 1997). The pCerulean-N1 Cyclin B1 (1147) was a gift from Jonathon Pines (Addgene plasmid #61852). The VSV-G (#12259), BFP and mCherry-NLS plasmids were obtained from Addgene.

### Induction of cell-cell fusion

Cell-cell fusion was induced following previous publications ((Feliciano *et al*., 2018). Briefly, cells were incubated for 20-30 seconds with a low pH buffer and immediate change into warm fresh medium or 4% PFA depending on the experimental set-up.

### Double synchronization assay for mitotic and G1/S or G2 cell-cell fusion

For G1/S + M cell-cell fusion, asynchronously growing LLC-PK1 cells were treated with 2 mM of thymidine, leading to a cell cycle arrest at the G1/S transition or S phase. After 14-18h, cells were released into fresh warm medium for 10h, after which a second 2mM thymidine treatment of 14-18h was applied together with 1.5 μM of nocodazole. With this set-up cells that during the release period were in G2 would become arrested in mitosis, due to the presence of nocodazole, and cells in G1 or S would be blocked at G1/S transition or in S phase by the presence of thymidine.

For G2 + M cell-cell fusion, asynchronously growing LLC-PK1 cells were incubated with 2mM thymidine for 14-18h. After this period cells were release into warm medium with 1.5 μM of nocodazole for 10h-12h, or until starting acquisition. Release from the thymidine block allowed cells to progress in the cell cycle and cells blocked in early or late S phase, by the thymidine treatment, would be expected to be, after 10h, in G2 or arrested in mitosis, respectively.

## Figure legends

**Figure S1 – Types of condensed DNA observed after induced mitotic entry.** Mitotic syncytium displaying three types of premature chromatin condensation: (1) duplicated mitotic chromosomes typical of G2 cells, (2) condensed single chromatids from early G1 cells and (3) unstructured DNA that from S phase cells. Grey shadow highlights the area of fused cells.

**Figure S2 – Histone H3 phosphorylation in anaphase control cells.** Analysis of pH3-s10 staining in control anaphase and telophase fixed LLC-PK1 cells, showing the progressive loss of phosphorylation from early anaphase to telophase.

**Figure S3 – LBR recruitment to the nuclear envelope after induced mitotic exit is defective.** (A) Time-lapse of EGFP-LBR recruitment to the nuclear envelope in control and mitotic cells forced to exit mitosis. EGFP-LBR was transiently transfected in LLC-PK1 cells expressing H2B-mCherry. Time is in h:min. Scale bar is 10μm. (B) Quantification of EGFP-LBR fluorescence intensity at the nucleus in control (n=2 cells) and mitotic cells with premature mitotic exit (n=3 cells). (C) Quantification of anaphase duration in control cells (n=5 cells) and time from fusion to nuclear envelope reformation in mitotic fused cells (n=7 cells).

**Figure S4 – Fusion of mitotic and interphase cells leads to aberrant syncytia.** Fusion of a mitotic cell with several interphase cells results, on one hand, in the formation of mitotic syncytium with defective spindles and become arrested in mitosis for several hours. On the other hand, in high ratios of interphase: mitotic cells, mitotic cells are forced to exit mitosis (interphase syncytium), reforming nuclear envelope membranes around condensed chromosomes.

## Notes

### Competing Interest Statement

The authors have declared no competing interest.

