## Supplementary figures and images for "Live dynamics of induced cell-cell fusion between mitotic and interphasic cells"

### Figure S1

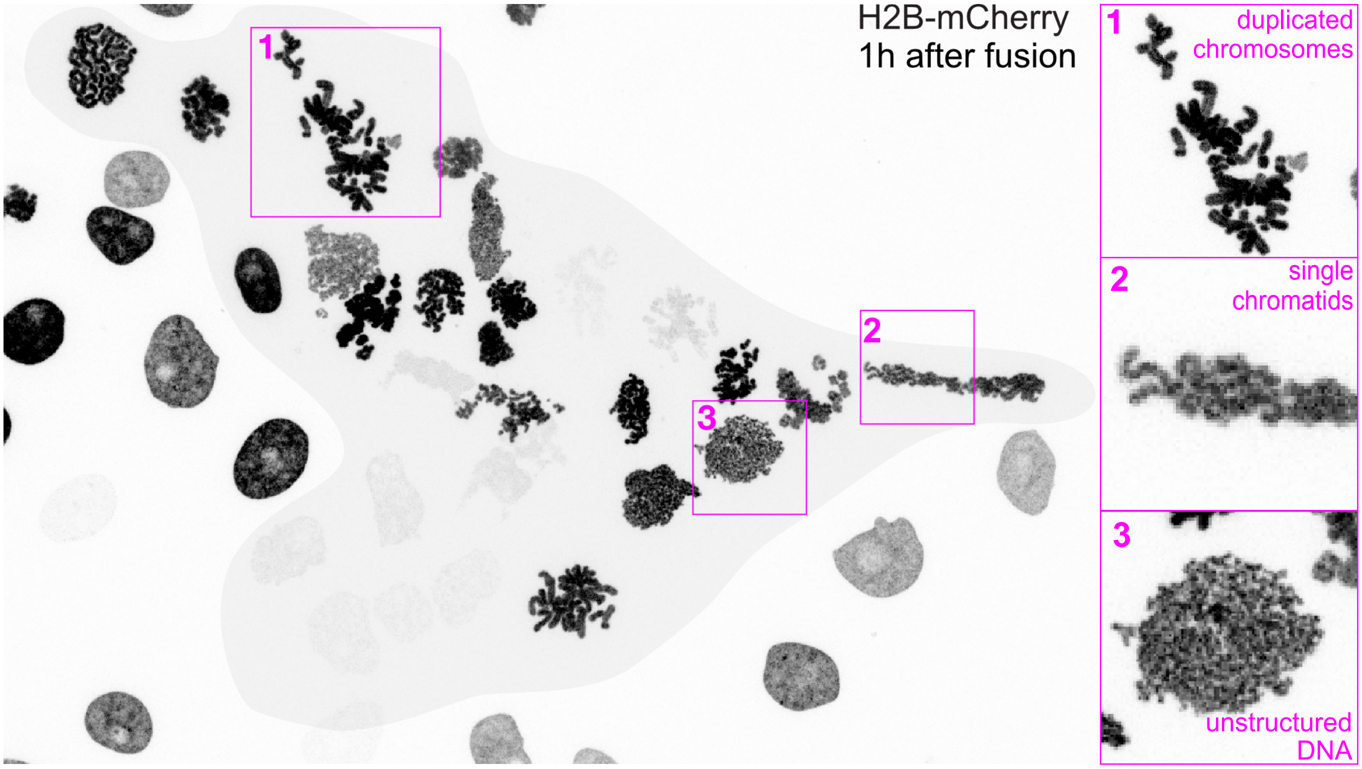

### Figure S2

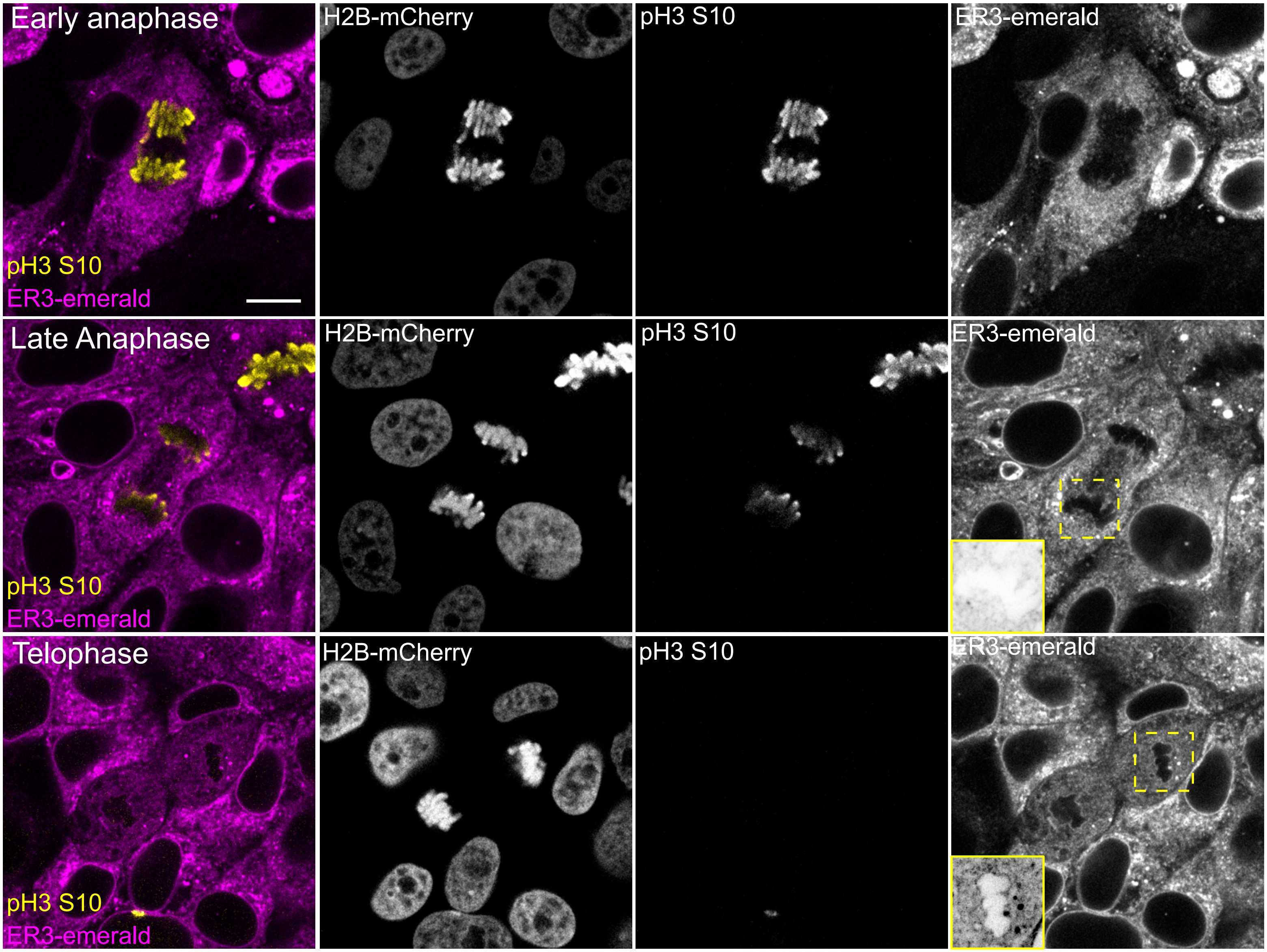

### Figure S4

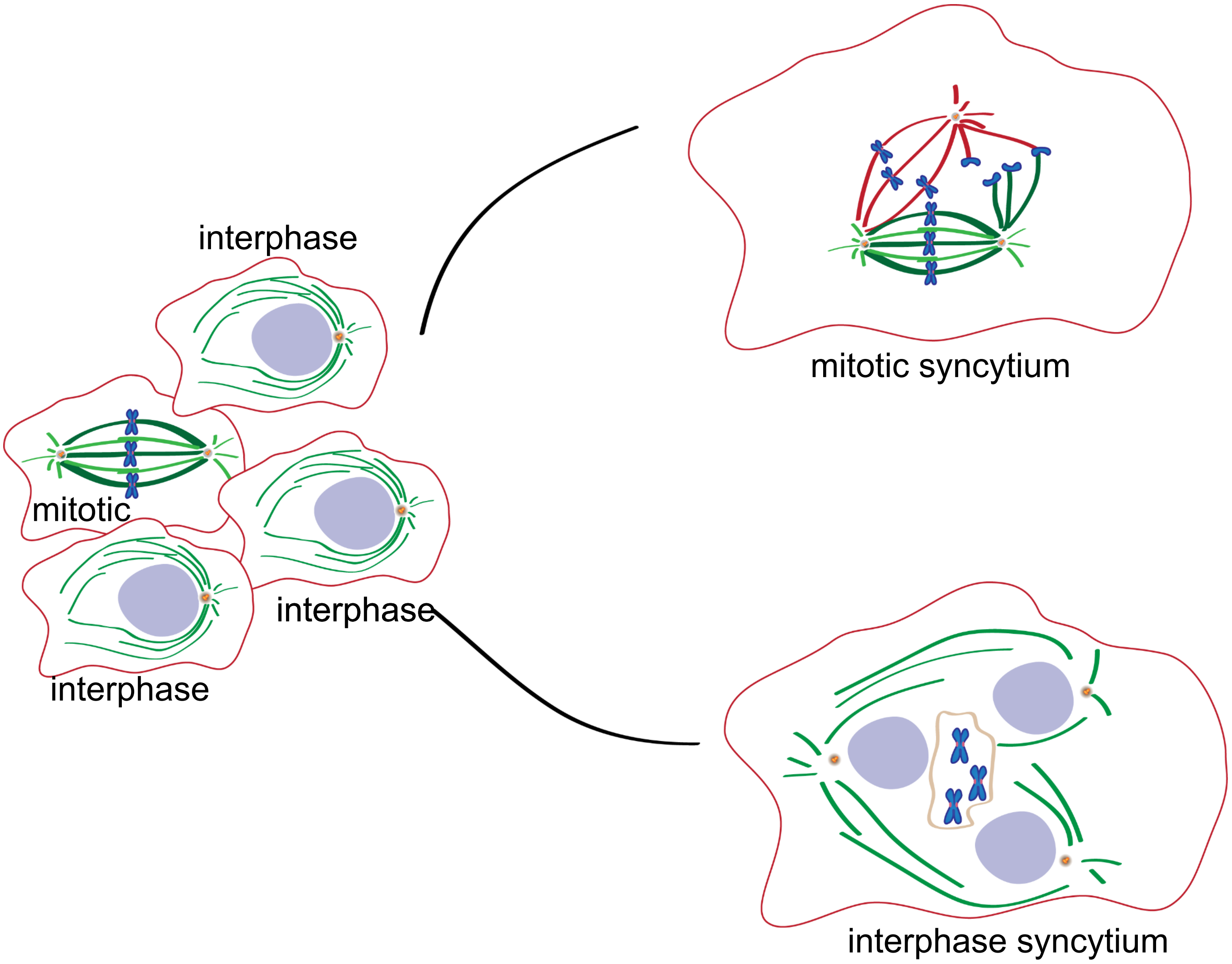

### Figures S3

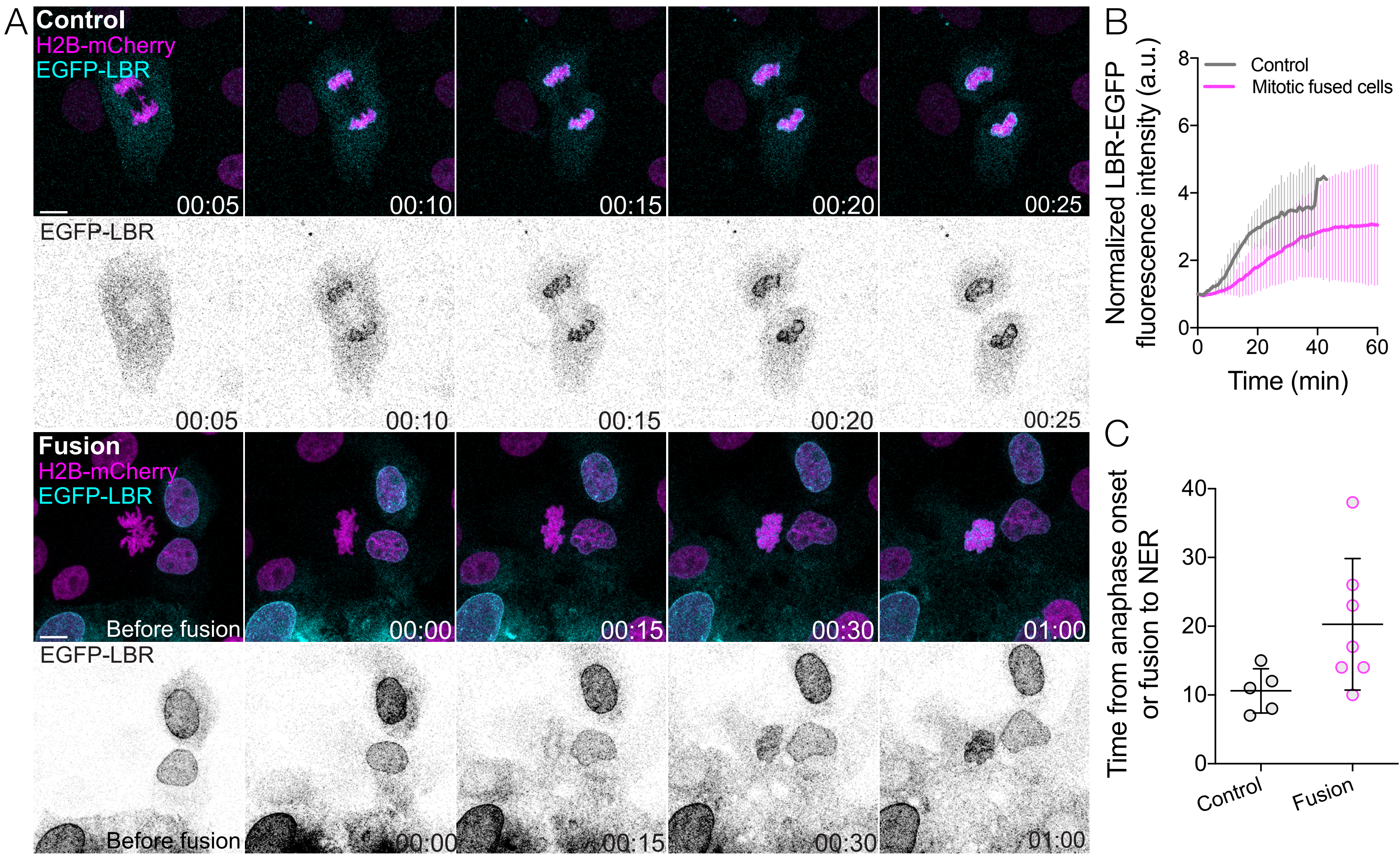
